# Sexually divergent DNA methylation programs with hippocampal aging

**DOI:** 10.1101/161752

**Authors:** Dustin R. Masser, Niran Hadad, Hunter Porter, Colleen A. Mangold, Archana Unnikrishnan, Matthew M. Ford, Cory B. Giles, Constantin Georgescu, Mikhail G. Dozmorov, Jonathan D. Wren, Arlan Richardson, David R. Stanford, Willard M. Freeman

## Abstract

DNA methylation is a central regulator of genome function and altered methylation patterns are indicative of biological aging and mortality. Age-related cellular, biochemical, and molecular changes in the hippocampus lead to cognitive impairments and greater vulnerability to neurodegenerative disease that varies between the sexes. The role of hippocampal epigenomic changes with aging in these processes is unknown as no genome-wide analyses of age-related methylation changes have considered the factor of sex in a controlled animal model. High-depth, genome-wide bisulfite sequencing of young (3 month) and old (24 month) male and female mouse hippocampus revealed that while total genomic methylation amounts did not change with aging, specific sites in CG and non-CG (CH) contexts demonstrated age-related increases or decreases in methylation that were predominantly sexually divergent. Differential methylation with age for both CG and CH sites was enriched in intergenic, and intronic regions and under-represented in promoters, CG islands and specific enhancer regions in both sexes suggesting that certain genomic elements are especially labile with aging, even if the exact genomic loci altered are predominantly sex-specific. Life-long sex differences in autosomal methylation at CG and CH sites were also observed. The lack of genome-wide hypomethylation, sexually divergent aging response, and autosomal sex differences at CG sites were confirmed in human data. These data reveal sex as a previously unappreciated central factor of hippocampal epigenomic changes with aging. In total, these data demonstrate an intricate regulation of DNA methylation with aging by sex, cytosine context, genomic location, and methylation level.

## Background

Direct DNA base modifications, such as cytosine methylation (mC), are proposed to fundamentally regulate the mammalian genome through altering genome accessibility (Law & Jacobsen 2010). Maladaptive changes in these epigenetic marks are potential drivers of pathogenesis and progression of many diseases (Robertson 2005). In the central nervous system (CNS), epigenetic changes have been associated with a number of age-related diseases, including Alzheimer’s, and cognitive impairment (Penner *et al.* 2010). Additionally, DNA modifications may regulate the gene expression program required for normal hippocampal learning and memory (Lister & Mukamel 2015). Despite the focus on age-related changes in mCG as a ‘biological clock’ (Horvath 2013) and the potential importance of DNA modifications in CNS aging (Lardenoije *et al.* 2015), DNA methylation changes in CG and CH (i.e., non-CG) contexts and genomic patterns with aging in the CNS of controlled animal models are largely unexplored. Additionally, studies have not examined the commonalities and differences between the sexes. The hippocampus is a central neural substrate of age-related dysfunction and disease but previous aging studies have not quantitatively examined mC genome-wide with single base resolution. Recent findings, albeit in other tissues, provide support for the need to examine alterations in DNA modification with aging (Hahn *et al.* 2017; Petkovich *et al.* 2017; Stubbs *et al.* 2017; Wang *et al.* 2017).

In an effort to elucidate the functional role of DNA methylation alterations with aging in the hippocampus, we recently reported that neither the expression of the major DNA methylation regulating enzymes (DNA methyltransferases and Ten-eleven translocases) nor the total mean genomic methylation or hydroxymethylation levels in either CG or CH contexts change with aging in the male or female hippocampus (Hadad *et al.* 2016). These results were unanticipated given the number of recent reviews that emphasize global DNA hypomethylation with age across somatic and CNS tissues as a driver of the aging process (Chow & Herrup 2015; Zampieri *et al.* 2015; Sen *et al.* 2016). A growing number of reports convincingly demonstrate that in the CNS mC can increase or decrease at specific CG (Penner *et al.* 2016; Ianov *et al.* 2017) and non-CG/CH (Mangold *et al.* 2017) sites and regions with aging. This suggests mechanisms whereby differential DNA methylation with aging is targeted in a locus-specific manner. Thus, quantitative analyses with single-base resolution are needed to identify where in the genome increases and decreases in methylation of cytosines are occurring with aging in the hippocampus and critically how sex factors into the epigenome response to aging.

This study sought to answer three questions: 1) Does global/total mean hippocampal methylation change with age; 2) what are the comparative responses of hippocampal methylation patterns with age in males and females; and 3) are there hippocampal life-long sex differences in autosomal methylation? Using published recommendations on terminology, (McCarthy *et al.* 2012) we define a sex difference as a difference between males and females that persists throughout the lifespan, and define sex divergence as a sex-specific response to a stimulus, such as aging (Figure S1).

DNA methylation was quantified in a base-specific manner across promoters, CG Islands and associated flanking regions, and gene regulatory regions from the hippocampus of male and female young (3 months) and old (24 months) C57BL6 mice using bisulfite oligonucleotide-capture sequencing (BOCS), a method we have quantitatively validated (Masser *et al.* 2016). Targeting 109Mb of the mouse genome (Hing *et al.* 2015)[over 30 million combined CG (∼3 Million) and CH (∼28 Million) sites] enables sequencing at depths exceeding coverage recommendations (Ziller *et al.* 2015) in multiple independent biological samples per group.

## Results

### Does global/total DNA methylation change with age?

DNA methylation was quantified in a base-specific manner (ie., methylation at each CG and CH was individually quantified) across almost all annotated promoters and CG Island units [Island, shore (±2 kb from island), and shelf (±2 kb from shores)] for a total of 109Mb of coverage using bisulfite-oligonucleotide capture sequencing (BOCS, Figure S2A-B). As the term CG island is often used non-specifically to refer to just the CG island or the combination of CG island and the accompanying shores and shelves, we will use CG island (CGI) to refer to the island alone, and a CGI unit will be used to refer to the island, shores, and shelves together. BOCS was developed to focus sequencing to regions of interest and avoiding repeat sequences, thereby increasing sequencing depth, while avoiding the bias towards CG dense regions observed and fundamental to reduced-representation bisulfite sequencing (Li *et al.* 2015; Masser *et al.* 2016). Over 30 million aligned reads were generated for each sample with a target enrichment of >25 fold from the whole genome for an average target sequence coverage of 20-40X (Figure S3). Using data from across all CG sites meeting sequencing coverage criteria sample-sample correlations were consistently very high (r>0.95), demonstrating robust reproducibility across all samples (Figure S4).

Mean levels of DNA methylation of all the CG and CH site calls in the targeted portion of the genome were calculated and there was no observed difference in global/total mean methylation levels across the hippocampal genome with age in males or females in either CG or CH contexts (Figure S5A and D). Furthermore, a consistent bimodal distribution of CG methylation was observed in both sexes at both ages (Figure S5B). No differences in the distribution of CG methylation in promoter and CGI units were evident as well (Figure S5C). Methylation at CH dinucleotides in the CNS has previously been shown to be higher relative to peripheral tissues (Lister *et al.* 2013; Lister & Mukamel 2015; Mangold *et al.* 2017), and similarly, we observe large numbers (>250,000) of individual CH sites (∼1% of all CH sites quantified) with high methylation levels (>10%) in the hippocampus (Figure S5E). While we have previously quantified the whole-genome levels of CH methylation in the hippocampus (Hadad *et al.* 2016), our previous approach did not yield base-specific data. This was also evident in the density distributions of CH sites subdivided into promoter regions or CGI units (Figure S5F). In total, there was no evidence for a genome-wide loss in methylation in either CG or CH sites, including when subdivided into promoter and CGI units.

### What are the comparative responses of mC to aging in males and females?

Next, whether specific sites in the genome were differentially methylated with aging in the female and male hippocampus was investigated. Age-related changes in mC were determined in a site-specific manner in males and females at individual CG (aDMCGs) and CH (aDMCHs) sites. aDMCGs were equally distributed between hypermethylation and hypomethylation events (Figure 1A, File S1). Comparing the aDMCGs in males and females, the majority (>90%) were sexually divergent (Figure 1B and File S1) though the extent of overlapping sites was greater than expected by chance (hypergeometric test, p<2.6E-106). This leaves the potential that different sites change in methylation level with aging in males and female but that these sites could be located in close proximity. A nearest neighbor analysis was performed between aDMCGs in males and females and the average distance was found to be >4kB between sex-specific aDMCGs in males and females. Differential methylation of CG sites with aging was observed across the genome (Figure 1C). Taking the union of all the aDMCGs in males and females samples were clustered and formed tight groupings based on age and sex (Figure 1D), further demonstrating the consistency of the methylation patterns. Animals separated by age in the 1^st^ component and by sex in the 2^nd^ component.

**Figure 1:**
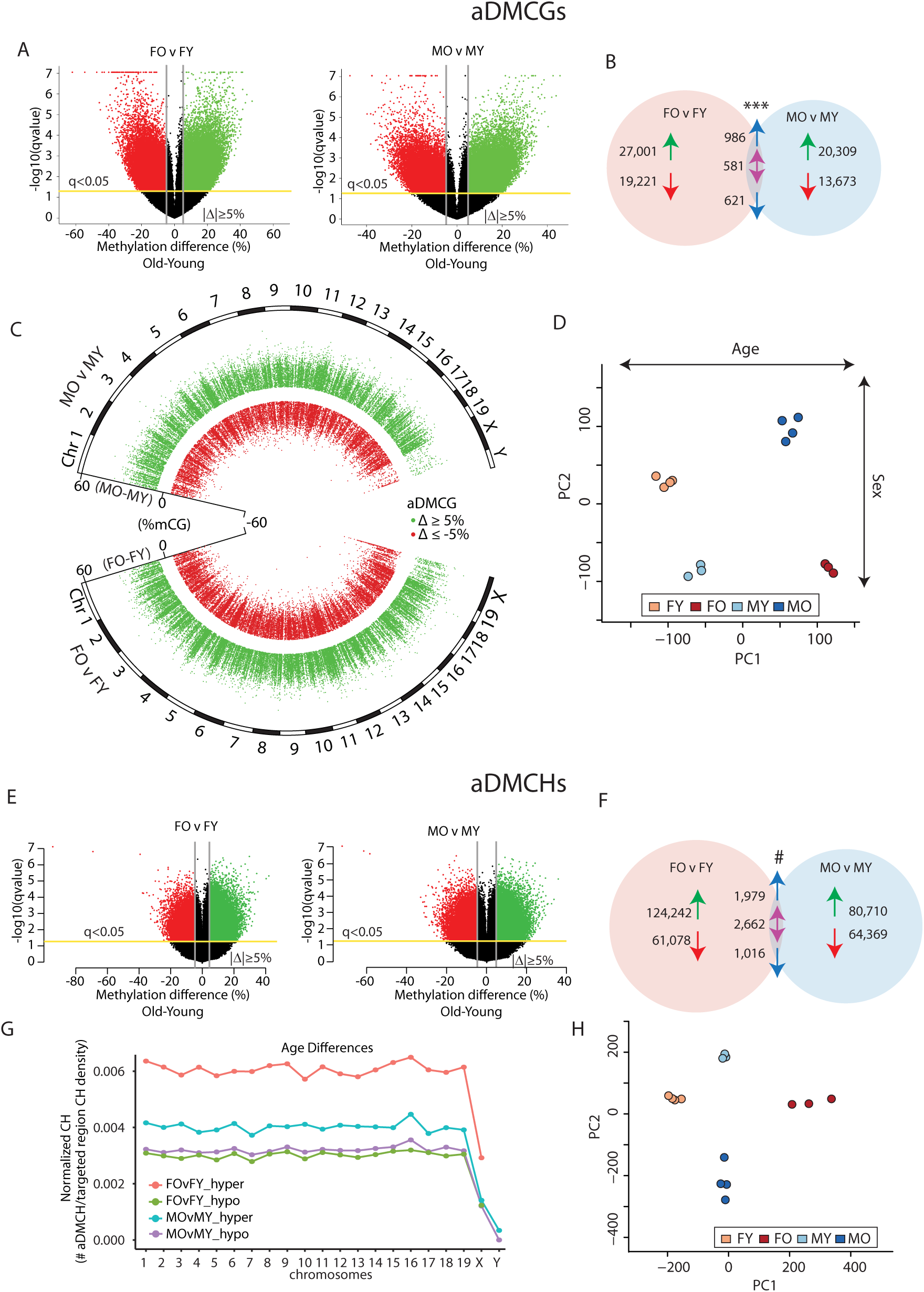
**Age-related differentially methylated CGs (aDMCG) and CHs (aDMCGH) in male and female hippocampus.** A) Volcano plots of pairwise comparisons of age-related CG sites methylation changes in females (Female Old – FO vs Female Young FY) and males (Male Old – MO vs Male Young MY). Sites with false discovery corrected p-value (q<0.05) and an absolute magnitude change Old-Young>|5%| were called differentially methylated. Sites with an ag-related increased methylation are represented in green and those with decreased methylation in red. B) Comparing the aDMCGs in males and females, more sites were found in common between the sexes than would be expected by chance (***, p<2.6E-106) but the majority of aDMCGs were sex specific. Arrows represent increased or decreased methylation with age and in the intersection blue arrows represent common regulation between the sexes and the double purple arrow represents different direction of change in the sexes with age. C) aDMCGs are presented by chromosomal location in males (top) and females (bottom) and the difference in mean methylation (Old-Young) on the inner axis. Each point represents one aDMCG meeting false discovery rate (FDR) cut off of q < 0.05. Sites ≥ 5 % absolute change in methylation with age (hypermethylated) are in green, while in red are sites ≤ −5% change with age (hypomethylated). D) The union of sites from B were used in a principle component analysis of the samples. Samples cluster by group and separated by age in the 1^st^ component and by sex in the 2^nd^ component. E) aDMCHs were compared in the same manner for differences in methylation with age in females and males. F) More sex common aDMCHs were observed than expected by chance (#, p<01E-200) but the majority of aDMCHs were specific to one sex or the other. Arrows as in B. G) Distribution of CH sites across the chromosomes examined by comparing the number of aDMCHs per the number of CHs in the covered regions of that chromosome. Equal representation of aDMCHs across the autosomes was observed with a lower rate of aDMCHs in the sex chromosomes. H) Taking the union of the sites in F, principle component analysis of the samples demonstrated tight clusters of samples by sex and age.

We have previously observed CNS tissue differences in MHCI promoter CH methylation with aging (Mangold *et al.* 2017), but there is little additional reported data on CH methylation with aging in the CNS (Lister *et al.* 2013). Age-related differentially methylated CHs (aDMCHs) were identified in a manner similar to aDMCGs. Hypermethylated and hypomethylated aDMCHs were observed with aging in males and females, with a greater number of hypermethylated sites (Figure 1E). Comparing the aDMCHs in males to females, the majority were sex specific but with greater overlap than would be expected by chance (hypergeometric test, p<01E-200) (Figure 1F). Unlike aDMCGs, aDMCHs in males and females are closer to one another as > 65% of these sites are within 1 kb of each other, likely due to the number of aDMCHs and the frequency of CHs in the genome compared to CGs. aDMCHs were found across the autosomes at a roughly equal density (Figure 1G). Clustering of samples by aDMCHs demonstrated close grouping of samples and separation of the groups by age and sex (Figure 1H).

In other organs, aging CG methylation changes demonstrate enrichment within common annotated genomic elements including CGIs, CGI shores, exons, and around transcriptional start sites (McClay *et al.* 2014; Bell *et al.* 2016). Sex common aDMCGs and sex specific aDMCGs in males and females were assessed for over-representation in genic (promoter, intron, exon), and CGI unit (island, shore, shelf) locations as compared to the distribution of CGs targeted in the oligonucleotide capture set for which there was coverage meeting our cutoffs. Enrichment of aDMCGs in outside of CGI units, in intergenic regions and in introns was evident (Figure 2A and B). CGIs and promoter regions were under-represented as were CGI shores, among hypermethylation events. Of note, while the specific regions or sites differentially methylated with aging are largely sex-specific, they demonstrate similar patterns of genic and CGI unit over-/under-representation in sex common and sex specific age-related changes regardless of whether the sites were hypomethylation or hypermethylation events. These results differ from those found in non-CNS tissue (McClay *et al.* 2014; Bell *et al.* 2016) and demonstrate non-CG dense intergenic and intronic regions as preferential locations for age-related hippocampal epigenetic alterations while CG dense regions like CGIs and promoters are under-represented in age-related changes in methylation. Additionally, there was not a significant difference in the localization of sex-common and sex-divergent differences with aging.

**Figure 2:**
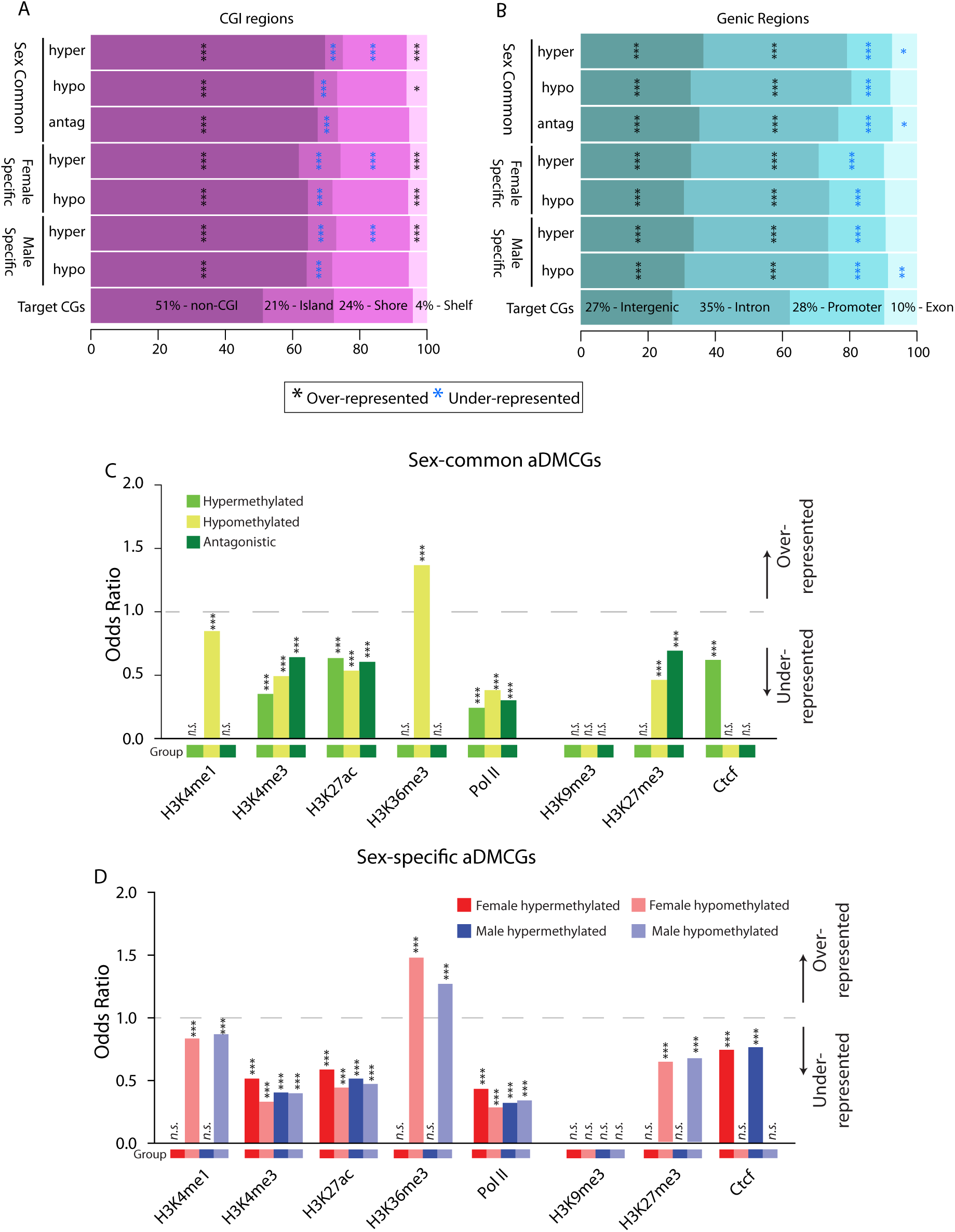
**Annotation enrichment patterns of age-related differentially methylated CG sites.** A) Sex common and sex specific aDMCG distributions were examined for enrichment in relation to the CG distribution in the CGI unit regions analyzed (target). aDMCGs were separated by whether they decreased or increased in methylation with aging or if they were antagonistically differentially methylated sex common sites (hypermethylated with aging in one sex and hypomethylated in the other sex). Over-representation of non-CGI regions and under-representation of Islands was observed for all comparisons. B) Similarly, genic regions were examined and aDMCGs were found to be over-represented in intergenic and intronic regions while under-represented in promoter regions. (***p<0.001, **p<.01, *p<0.05 χ^2^ analysis, coloring by over-representation, black or under-representation, blue.) C) Odds ratios demonstrating enrichment of sex-common aging differentially methylated CG sites (aDMCGs) for ENCODE and regulatory elements (activation - H3K4me1, H3K4me3, H3K27ac, H3K36me3, PolII, and repression - H3K9me3, H3K27me3, Ctcf) by GenomeRunner analysis. Enrichment comparisons were carried out for hypermethylated aDMCGs (light green), hypomethylated aDMCGs (yellow), and antagonistic (dark green) sex-common aDMCGs. Odds Ratios greater than 1.0 (gray dotted line) demonstrate over-represented while those less than 1.0 are under-represented. Significant enrichment or depletion are denoted by stars where * p < 0.05, ** p < 0.01, and *** p < 0.001. D) Odds ratios for sex-specific aDMCGs enrichment for ENCODE and regulatory elements. Enrichment comparisons were carried out in each sex for hypermethylated aDMCGs (dark red – females, dark blue - males) and hypomethylated aDMCGs (light red – females, light blue - males). Odds Ratios greater than 1.0 (gray dotted line) are over-represented while those less than 1.0 are under-represented. Significant enrichment or depletion are denoted by stars where * p < 0.05, ** p < 0.01, and *** p < 0.001.

Enrichment of aDMCGs in enhancer regions was also performed against ENCODE datasets of mouse brain tissue enhancer locations using GenomeRunner (Dozmorov *et al.* 2012) (Figure 2C and 2D). Regions associated with active transcription, H3K4me3, H3K27ac, and PolII were generally under-represented as a location for aDMCGs, regardless of the whether the site was hyper-or hypo-methylated with aging. Sex common and sex specific hypomethylated sites were enriched at the exonically localized H3K36me3 and under-represented at the primed enhancer marker H3K4me1. The repressive mark H2K27me3 was under-represented in hypomethylation aDMCGs while sex common hypomethylation events and sex specific hypermethylated sites were under-represented in Ctcf regions. Sex specific aDMCGs demonstrated similar patterns of enhancer enrichment in males and females, though with slight differences in the odds ratios (Figure 2D).

aDMCHs demonstrated similar but not identical patterns of enrichment for genomic elements. aDMCHs were under-represented in CGIs (with the exception of hypermethylation aDMCHs in females) and shores while enriched in non-CGI unit regions (Figure 3A). In genic regions aDMCHs were under-represented in promoter and exonic elements while enriched in intergenic, and in most comparisons intronic regions (Figure 3B). Much like sex common aDMCGs, sex common aDMCHs were significantly under-represented among activation associated enhancer elements H3K4me3, H3K27ac, and PolII and the repressive marks H3K9me3 and Ctcf (Figure 3C). However, unlike aDMCGs, sex-specific aDMCHs were over-represented at active H3K4me1 marks and no enrichment was found at H3K36me3 associated regions (Figure 3D).

**Figure 3:**
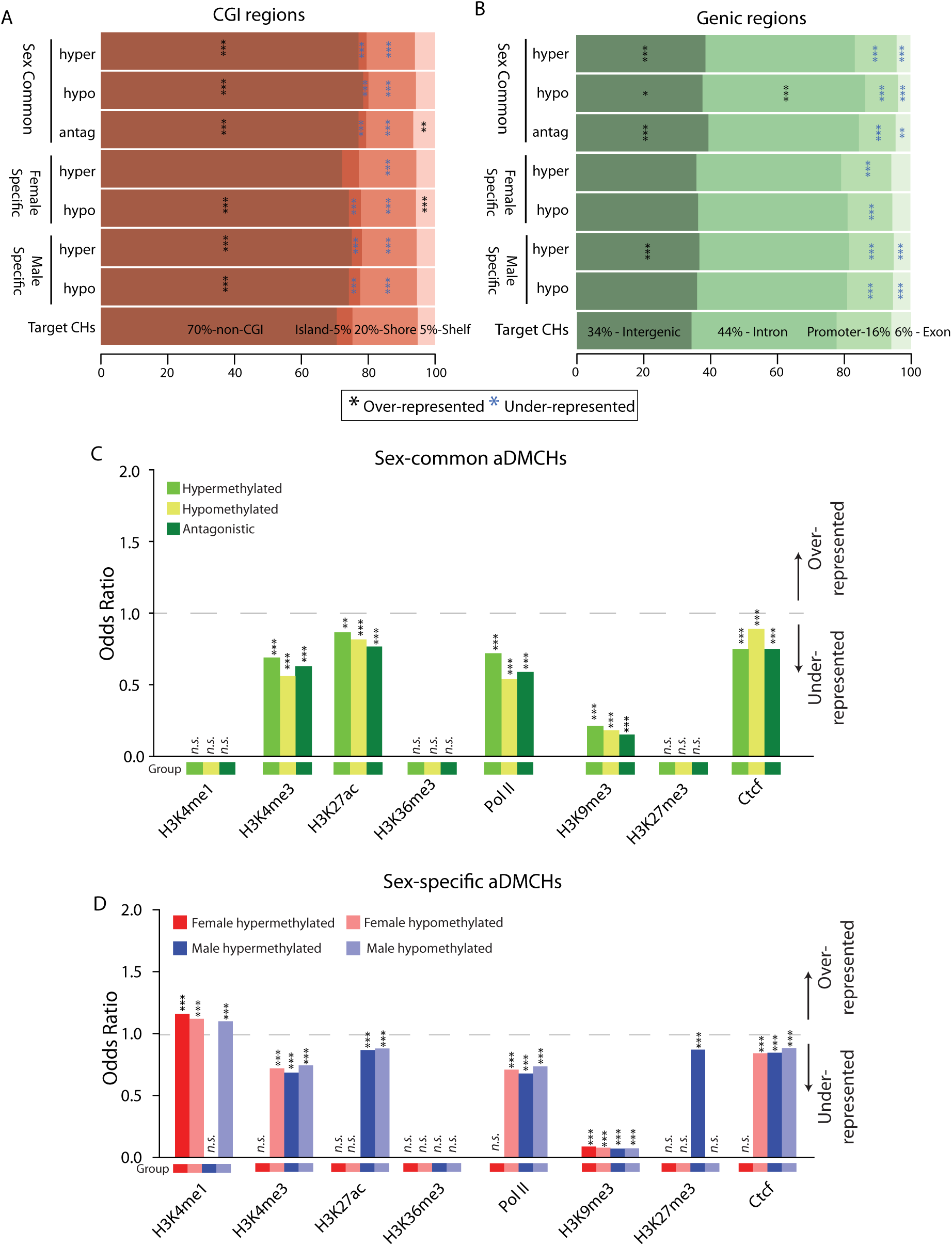
**Annotation enrichment patterns of age-related differentially methylated CH sites.** A) aDMCH distributions for males and females in the CGI unit regions analyzed (target) demonstrated over-representation in non-CGI regions and under-representation in CGI shores and islands themselves (with one exception). In many cases CGI shelves were over-represented for aDMCHs. B) In genic regions, aDMCHs were found to be over-represented in intergenic and under-represented in promoter and exonic regions. For a number of groups, especially hypomethylation, introns were over-represented as well. (***p<0.001, **p<.01, *p<0.05 χ^2^ analysis, coloring by over-representation, black or under-representation, blue.) C) Odds ratios of sex-common aDMCHs for ENCODE and regulatory elements by GenomeRunner analysis. Enrichment comparisons were carried out for hypermethylated aDMCHs (light green), hypomethylated aDMCHs (yellow), and antagonistic (dark green) sex-common aDMCHs. Odds Ratios greater than 1.0 (gray dotted line) are over-represented while those less than 1.0 are under-represented. Significant enrichment or depletion are denoted by stars where * p < 0.05, ** p < 0.01, and *** p < 0.001. D) Odds ratios of sex-specific aDMCHs enrichment for ENCODE and regulatory elements. Enrichment comparisons were carried out by sex for hypermethylated aDMCHs (dark red – females, dark blue - males) and hypomethylated aDMCHs (light red – females, light blue - males). Odds Ratios greater than 1.0 (gray dotted line) are over-represented while those less than 1.0 are under-represented. Significant enrichment or depletion are denoted by stars where * p < 0.05, ** p < 0.01, and *** p < 0.001.

### Are there life-long sex differences in autosomal methylation?

While sex differences in methylation patterns are evident during development (Nugent *et al.* 2015), little is known about whether brain sex differences persist throughout life. Sex differences in methylation of 901 CG (sDMCG) and 3,028 CH (sDMCH) sites were found at both young and old ages (e.g., higher in males than females at both young and old age, or vice versa). Analysis of sex differences was restricted to autosomes and sex differences were found throughout the autosomes for both CGs (Figure 4A) and CHs (Figure 4B, File S2). The difference in methylation between males and females was also found to be stable between young and old ages at these sites. These life-long sex differences in CG and CH site methylation were significantly enriched in intergenic, intronic, and non-CGI unit regions while generally under-represented in promoters, exons, CGIs, shores, with some exceptions (Figure 4C and D). Additionally, CH sites that were higher in males were significantly enriched in introns, while the CH sites with higher methylation in females were not (Figure 4D). For both sDMCGs (Figure 4E) and sDMCHs (Figure 4F) sex differences were not enriched for either active or repressive enhancer elements. These findings demonstrate that while not extensive, sex differences in hippocampal CG and CH methylation are evident in early adulthood (3M) and persist into advanced age.

**Figure 4:**
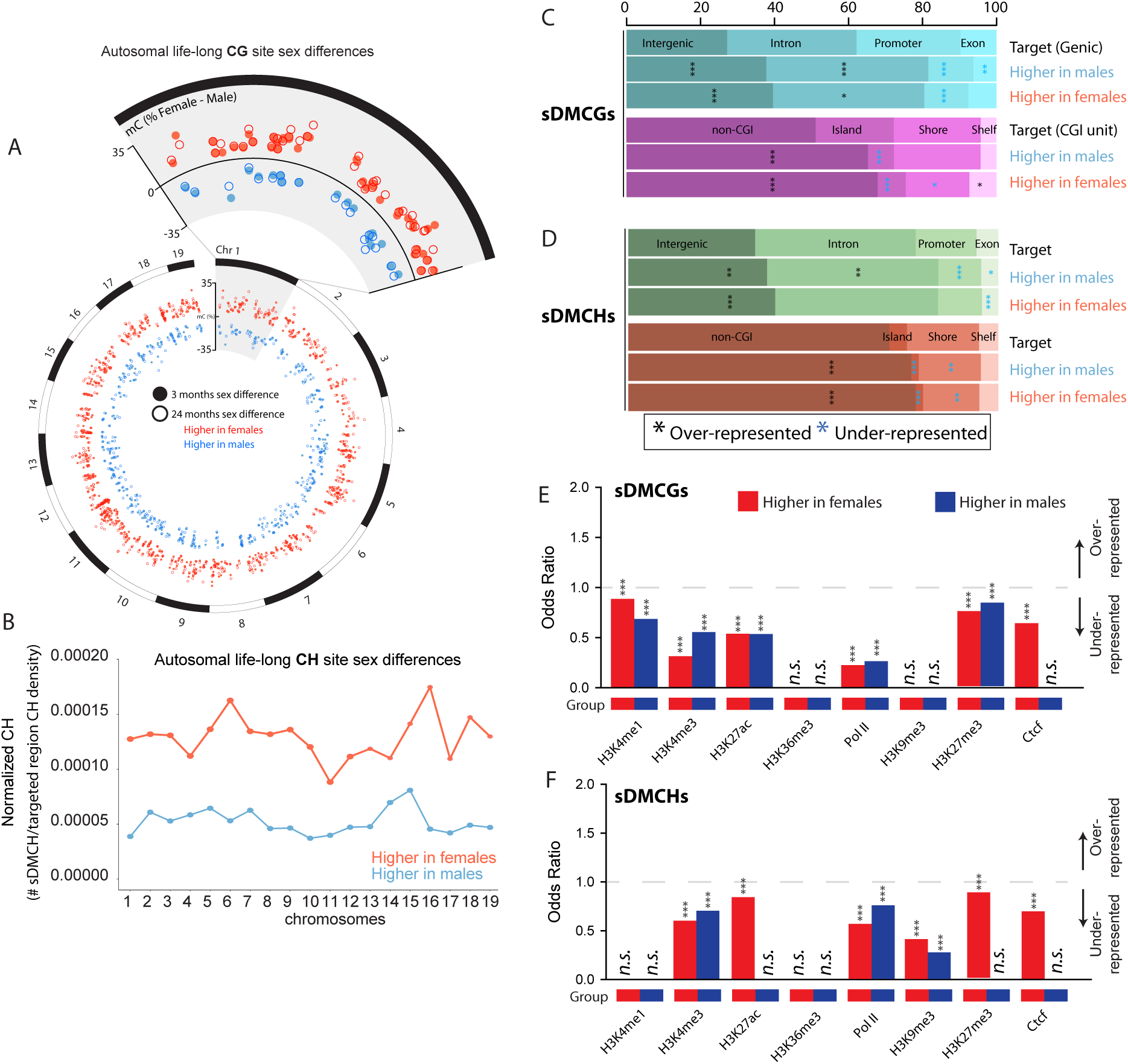
**Life-long sex differences in autosomal DNA methylation.** A) Autosomal locations of the 901 life-long CG site sex differences (sDMCGs) with higher methylation levels in females (orange) and sites that have higher methylation in males (blue). Closed circles represent the methylation difference between sexes in young (3 months) mice, while open circles represent the methylation difference between sexes in old (24 months) mice. For clarity, chromosome 1 is enlarged to visualize the life-long nature of these autosomal sex differences. B) Autosomal distribution of life-long CH site sex differences (sDMCHs) for the 3,028 CH sites that have either higher methylation levels in females (orange) or higher methylation in males (blue) relative to the CH density of the region examined. C) sDMCG and D) sDMCH site enrichment profiles among Genic (top) and CGI unit (bottom) regions for sites with higher methylation levels in males or in females. Percentages of sDMCGs and sDMCHs in each region type are presented in Figures 2 and 3 (***p<0.001, **p<.01, *p<0.05 χ^2^ analysis, coloring by over-representation, black or under-representation, blue). E) Odds ratios of sDMCG and F) sDMCH enrichment in ENCODE and regulatory elements by GenomeRunner analysis. Odds Ratios greater than 1.0 (gray dotted line) are over-represented while those less than 1.0 are under-represented. Significant enrichment or depletion are denoted by stars where * p < 0.05, ** p < 0.01, and *** p < 0.001.

### Replication in humans

To validate the primary findings of the mouse studies, namely lack of hypomethylation with age, sex-common and sex divergent differences with aging, and life-long sex differences; we collected publicly available datasets from control human brain samples across a range of ages. Data from 19 hippocampal and 145 frontal cortex control (non-diseased) samples aged 13-95 were collected. These data were generated with methylation microarrays and provide CG methylation quantitation across ∼450,000 probes. When assaying hypomethylation with age, a linear model of age’s effects on mean methylation of any sample (methylation ∼ age) showed neither a significant effect in hippocampus (p = .228, R^2^ = .07185) nor in frontal cortex (p = .543, R^2^ = .002587) (Figure 5A). As sufficient sample size in males and females was not available for hippocampus, frontal cortex data (69 females and 70 males) was used to determine specific differential methylation with aging. Sites were analyzed for the factors of sex and age and for interaction effects (methylation ∼ age, methylation ∼ sex, methylation ∼ sex:age). A majority of sites (22,204) demonstrated a main effect of age alone, 2,723 were found to have a main effect of sex alone, and 4,154 sites showed both sex and age effects with a significant interaction. The sites with only an aging effect were attributed to sex-common age-related changes while sites with only an effect of sex were sex differences. A significant interaction effect was indicative of sites with age-related sex divergences. As presented in Figure 5B, these effects were generally evenly distributed – increases and decreases in methylation with age and sites with higher methylation in males or in females. Examining examples of each phenotype clear sex common age-changes (Figure 5C), sex divergences with aging (Figure 5D), and autosomal (sites were limited to only autosomes) life-long sex differences (Figure 5E) were observed. These human data collected and analyzed by different methods than the mouse data presented here demonstrate that the general principles of the mouse findings also occur with aging in the human CNS.

**Figure 5:**
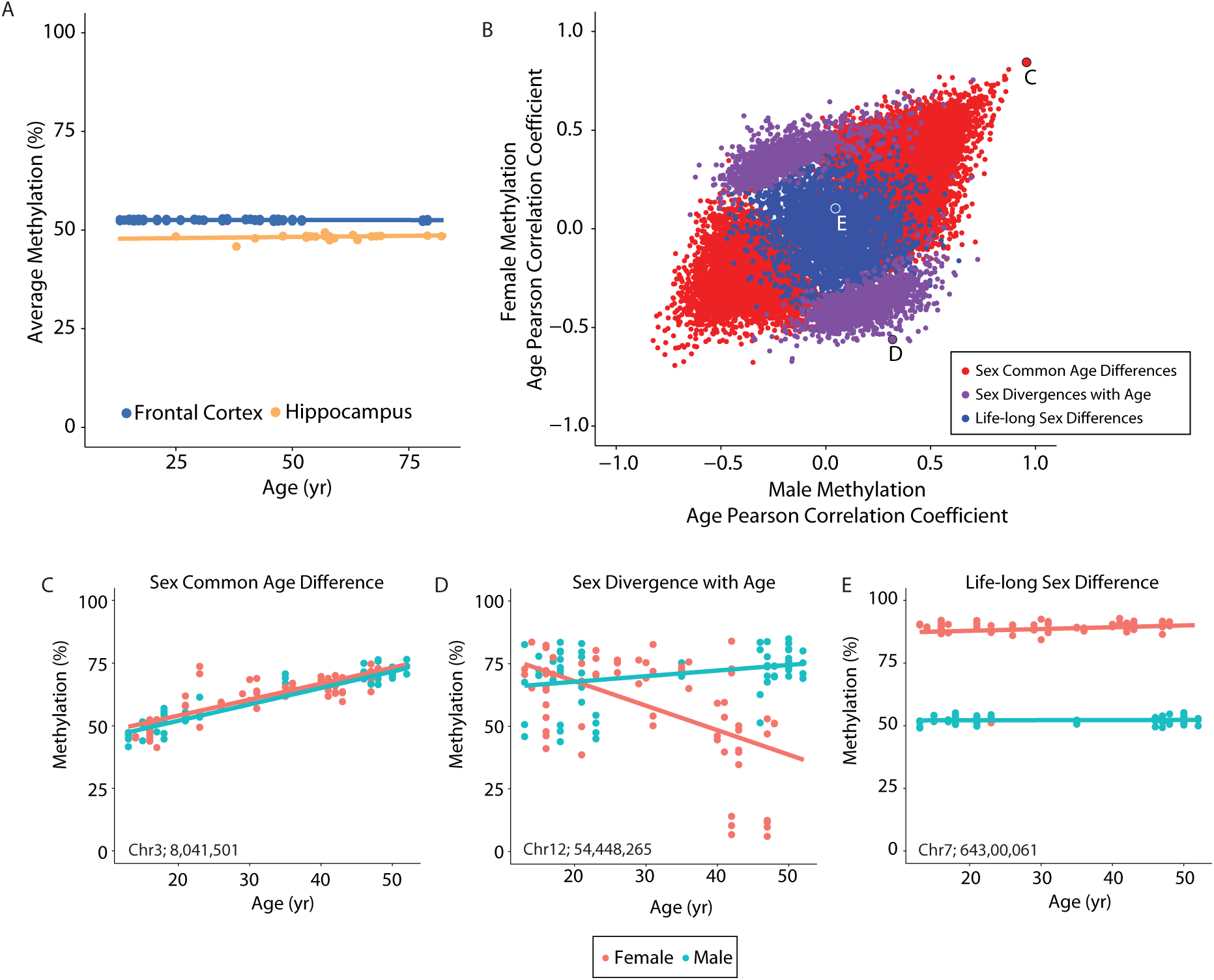
**Methylation changes with aging and sex differences in the human CNS.** A) Publicly available human methylation data from hippocampus and frontal cortex demonstrate no change in mean CG methylation with age. B) Using the fontal cortex data, for which there is a larger samples size and equal distribution between sexes, a general linear model was used to determine individual sites with significant age, sex or interaction effects on methylation level. Plotted by Pearson correlation coefficients to age by females (y-axis) and males (x-axis), sites with sex common age-related decreases in methylation (red, bottom left) and increases in methylation (red, top left) are evident. Sites with sexually divergent response to aging (significant interaction effect) are in purple. Life-long sex differences are plotted in blue. Sites without any significant factor of age or sex are not shown to improve clarity. C) Example site of a sex common age-related differentially methylated site. D) Example of sexually divergent response to aging. E) Example of life-long sex difference. Locations of specific sites are given and are also highlighted in panel B.

## Discussion

This is the first comprehensive genome-wide and base-specific quantitation study of mC patterns and alterations with age in the male and female hippocampus. A delineation of the patterns of age-related changes in DNA methylation is necessary for understanding both the potential functional effects of these changes and the regulation of DNA methylation patterns with aging. This study demonstrates a number of important principles for hippocampal DNA methylation: 1) genome-wide hypomethylation with aging does not occur; 2) site-specific differential methylation of both CG and CH sites occurs with aging; 3) males and females share some sites of differential methylation but age-related changes are primarily sexually divergent; 4) life-long autosomal sex-differences in CG and CH methylation are evident; and 5) these findings are replicated in human CNS data.

### Genome-wide hypomethylation with aging

No decreases in genome-wide CG or CH methylation were evident with aging in this study in either mice or humans. It is often stated that DNA methylation decreases globally with age across tissues (Ashapkin *et al.* 2015; Chow & Herrup 2015; Xu 2015; Zampieri *et al.* 2015; Sen *et al.* 2016) on the basis of reports using older chromatographic methods with low replicate numbers and no statistical analyses (Wilson & Jones 1983; Singhal *et al.* 1987). With modern sequencing tools this long-standing hypothesis of an age-related loss in global methylation can be revisited with more quantitatively accurate and validated tools. Taken with our previous findings of no changes in mean hippocampal DNA methylation with aging as determined by low coverage whole-genome oxidative bisulfite sequencing and pyrosequencing of LINE and SINE repeat elements (Hadad *et al.* 2016) genomic hypomethylation with aging is not evident in the hippocampus. Studies published since the submission of this report examined liver, lung, heart, and cortex using bisulfite sequencing methods have also found no genome-wide hypomethylation with aging (Cole *et al.* 2017; Stubbs *et al.* 2017; Wang *et al.* 2017).

### CG and CH methylation are regulated with aging

A principle finding from this study is that CH methylation changes in a base-specific fashion between adulthood and old age. Prior brain aging and methylation studies have not examined CH, also referred to as non-CpG, methylation. Large numbers of CH sites were differentially methylated with aging, over 100,000 in both males and females. Widespread methylation (>10% mC) of CH sites agrees with previous findings that the CNS contains some of the highest mCH levels in the body (Lister *et al.* 2013). CG dinucleotides are under-represented in the mammalian genome, while CH contexts are much more common (∼30 fold more). This is apparent in our BOCS target regions as they contain over 28 million cytosines in the CH context, and approximately 3 million in the CG context. The reproducible methylation of specific CH sites we observed here and have previously reported (Mangold *et al.* 2017) argues against accidental or non-regulated methylation of these sites. The functional role of CH methylation in the mammalian genome is only beginning to be understood and requires further testing (He & Ecker 2015). Given the important developmental role of mCH in synapse development, dysregulation of mCH with aging could have important functional impacts (Lister *et al.* 2013).

Additionally, as mCH has differences in writing, erasing, and reading mechanisms as compared to mCG, altered methylation at CH sites could have functionally distinct impacts from CG methylation changes (Kinde *et al.* 2015; Mo *et al.* 2015). These data provide the first view, to our knowledge, that specific CH sites across the genome are differentially methylated with aging in the hippocampus. When possible, such as with bisulfite sequencing approaches, inclusion of mCH analysis in brain aging studies is warranted and future cell type specific studies will be able to determine if aDMCHs are preferentially occurring in neurons, as neurons have higher levels of mCH than non-neuronal cell types in the brain (Lister *et al.* 2013).

### Sex divergence of differential methylation with aging

Prior studies of age-related changes in brain cytosine methylation have generally not examined the factor of sex in the patterns of changes. While there are sex divergences in methylation during development (Nugent *et al.* 2015) and we have reported sex divergences with aging in MHCI promoters (Mangold *et al.* 2017), the base-specific differences in methylation changes with aging across the genome in both sexes have not been examined [see Figure S1 for graphical definitions of sex divergence and difference according to (McCarthy *et al.* 2012)]. While males and females share many more sex common age-related changes than would be expected by chance, the majority of aDMCGs and aDMCHs were sexually divergent. Our findings reveal a previously unappreciated central factor of sex in age-related epigenetic changes. Importantly, this sexual divergence was not simply the result of a loss in sex differences with aging, which were observed to persist throughout life as discussed below. A recent report (Stubbs *et al.* 2017) found that a mouse methylation ‘age’ was similar in males and females and that sex hormones influence methylation changes with aging. Taken together with our results it is clear that the epigenomic response to aging is a combination of sites regulated in both sexes and sites regulated in only one sex. This was confirmed in human data as well. As discussed below the enrichment of aDMCGs and aDMCHs in annotated genomic regions and enhancers is quite similar providing evidence that these sex divergences are not generally occurring in different types of genomic locations, rather there is some form of spatial regulation by sex on the exact location of aDMCGs and aDMCHs. Future studies of epigenomic changes with brain aging will need to incorporate both males and females into study designs and sex-specific databases of enhancer locations need to be developed to improve interpretation of this data.

### Enrichment of age-related changes to annotated genomic regions

Recent studies in liver and other tissues examining age-related methylation changes find limited agreement in the types of genomic regions (*e.g.* introns, exons, promoters) where differences are likely or unlikely to occur (Cole *et al.* 2017; Hahn *et al.* 2017; Stubbs *et al.* 2017), and aDMCGs observed here share commonalities and differences in enriched regions with these studies. Therefore, we focused on comparing the genomic enrichment between aDMCGs and aDMCHs, males and females, and between hypermethylated and hypomethylated sites. Enhancers have been identified previously to be genomic regions with dynamic DNA methylation with aging (Jones *et al.* 2015). For example, the presence of H3K27ac and H3K4me1 marks are indicative of active enhancer regions, while H3K4me3 marks are enriched in active promoters (Chen & Dent 2014). The presence or absence of H3K27ac at enhancer regions distinguishes enhancers as active or inactive/poised, respectively (Creyghton *et al.* 2010). Interestingly, H3K36me3 is suggested to define exon boundaries and play a role in exon selection during transcription (Schwartz *et al.* 2009). Together the relative enrichment or depletion of these different genomic regions demonstrates that the brain has a unique pattern of regions where age-related methylation changes are likely or unlikely to occur compared to other tissues. It should be noted that ENCODE datasets were generated against young male mice (Mouse ENCODE consotium *et al.* 2012). Age -and sex-specific enhancer annotations as well additional data on chromatin structure will aid in placing methylation changes in their appropriate context (Pal & Tyler 2016). This is emphasized by the finding that the genomic regions enriched in age-related methylation changes differ from other tissues.

### Life-long autosomal sex differences

While more limited in number than age-related differences, life-long autosomal CG and CH methylation is evident in the mouse hippocampus. Developmental sex divergences in CNS methylation have been recently reported (Nugent *et al.* 2015). Here we observe an additional phenomenon – differences between males and females that are evident after development and are stable throughout life. While differences in sex chromosome methylation and X and Y encoded transcript expression are well known these data demonstrate a further level of autosomal sex differences. This was found to be true in both mice and humans. The functional implications of life-long autosome sex differences remain to be determined and sex differences in CH methylation have, to our knowledge, not been previously reported. sDMCGs and sDMCHs were enriched in intergenic, intronic and non-CGI unit regions while under-represented in CG islands and most of the enhancer regions examined. As stated previously, integration of this data with enhancer elements remains difficult due to the lack of sex-specific enhancer data for comparison and is an area of needed study.

### Future studies

These findings raise important questions to be addressed in future studies. The epigenomic regulatory processes that could lead to the profound sex divergence with aging observed are unknown. While epigenetic mechanisms establishing methylation patterns during organismal development have been extensively studied, the mechanisms whereby methylation or de-methylation is directed to specific genomic locations with aging are not known. Potential mechanisms include hormonal modulation of DNMT activity, which may explain sex-differences with aging and sexual divergent epigenetic responses to aging, but the signal gating that targets changes to specific loci require extensive further investigation. The sexual divergence with aging described here could potentially be harnessed to help understand these regulatory processes as these differences are naturally occurring and not the result of a genetic intervention.

While this study did not distinguish between mC and hmC, future studies will need to identify how hydroxymethylation changes within and between sexes with aging using oxidative bisulfite sequencing (Hadad *et al.* 2016). Analysis of isolated, specific CNS cell populations (e.g., microglia, neurons) should also be an area for further investigation. As well, analysis of a range of ages across the lifespan would enable determination of whether these age-related changes slowly accumulate with time or if there are periods in the lifespan with more abrupt changes in genomic methylation patterns.

In summary, our results present novel evidence for sexual divergent DNA methylation programs with hippocampal aging in patterns of CG and CH methylation, life-long sex differences in CG and CH methylation and confirm the CG findings in human methylation data. The NIH recommends inclusion of animals of both sexes in studies when warranted and these results provide a clear rationale for the need to examine both males and females and perform single base resolution analysis of DNA methylation patterns. Furthermore, our data highlight the complexity of the regulation and functional significance of epigenetics with aging, specifically DNA methylation, in the CNS.

## Experimental Procedures

Detailed descriptions of experimental procedures, reagents, and associated references can be found in online supporting information.

### Animals

All animal experiments were performed according to protocols approved by the Penn State University Institutional Animal Care and Use Committee. Male (N=8, n= 4 young and n=4 old) and female (N=8, n=4 young and n=4 old) C57BL6 mice ages 3 (young) and 24 (old) months were purchased from the National Institute on Aging colony at Charles River Laboratories (Wilmington, MA). Mice were housed in the specific pathogen-free Pennsylvania State University College of Medicine Hershey Center for Applied Research facility in ventilated HEPA filtered cages with *ad libitum* access to sterile chow (Harlan 2918 irradiated diet, Indianapolis, IN) and water. While in the facility all animals were free of helicobacter and parvovirus. Following a one week acclimation period on entering the respective facility, male mice were euthanized by decapitation. Female mice were euthanized by decapitation during diestrus after estrous cycle staging.

## Acknowledgements

The authors thank Drs. Graham Wiley and Patrick Gaffney of the OMRF Genomics facility, the OU Supercomputing Center for Education and Research, and A. Nathaniel Joiner for assistance with figure generation. This work was supported the Donald W. Reynolds Foundation, the Oklahoma Nathan Shock Center of Excellence in the Biology of Aging Targeted DNA Methylation and Mitochondrial Heteroplasmy Core (P30AG050911), the National Institute on Aging (R01AG026607, F31AG038285, T32AG052363), National Eye Institute (R01EY021716, R21EY024520, T32EY023202) and Oklahoma Center for Advancement of Science and Technology (HR14-174).

## Authors’ contributions

DRM, CAM, AR, and WMF designed and executed the study. DRM, NH, AU, CBG, MGD, JDW, DRS, and WMF analyzed the data. MMF provided animal support. All authors read and approved this manuscript.

## Supporting Information Listing

**Supplemental Figure 1:**
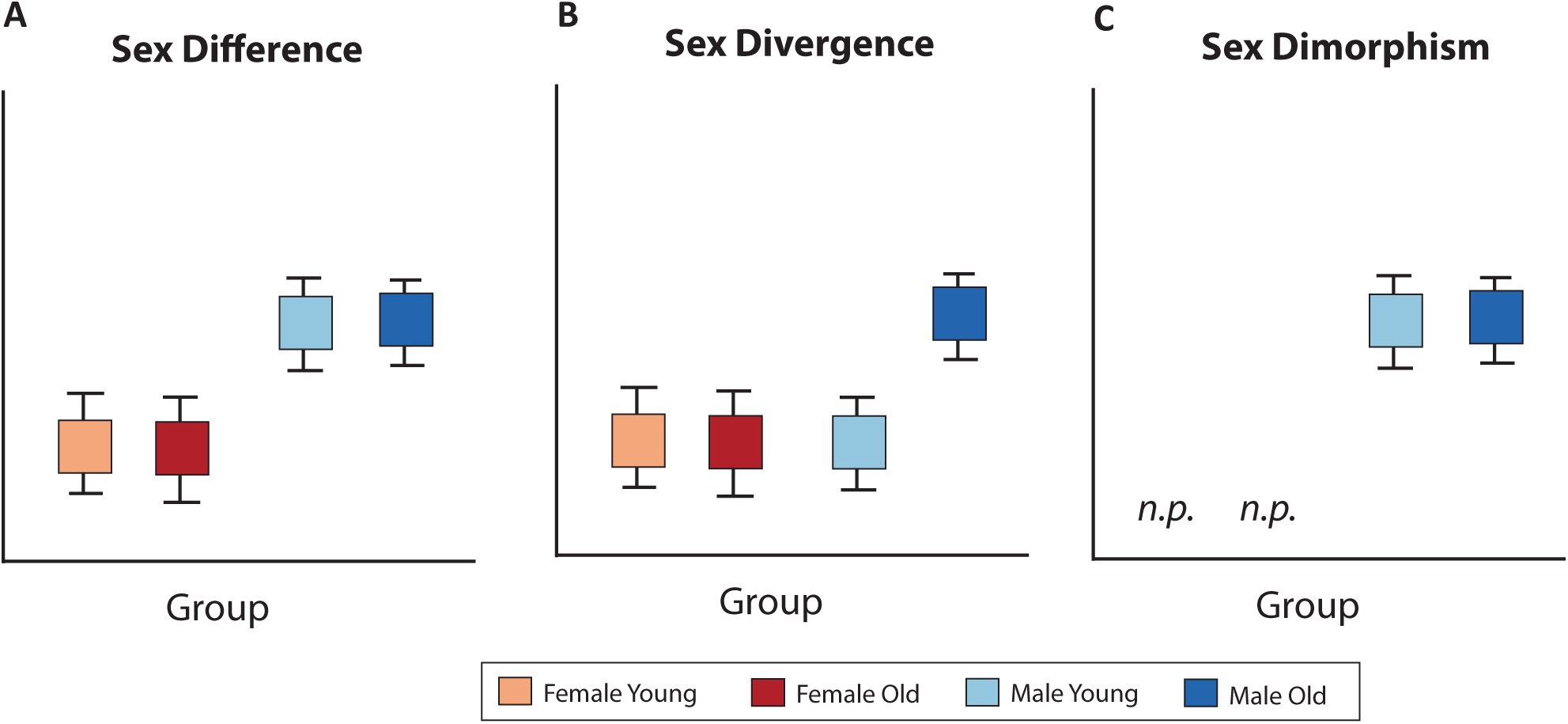
**Terminology definitions** A) Sex differences are defined as a difference in the mean between males and females and for the purposes of this study independant of age, i.e. life-long. B) Sex divergence is a difference in response to aging that is only present in one sex, in this example an increase in males only with aging. Sex divergences can occur in either sex and can be increases or decreases. C) Sex dimorphisms are a dialectic difference between males and females where the endpoint is present in one sex, in this case males, and absent (not present, *n.p.*) in the other sex. Examples of sex dimorphisms include sex organs which are present in only one sex. Sex dimor-phisms are not relavent for the present study as the cytosines examined are present in both males and females, only the level of methylation is different.

**Supplemental Figure 2:**
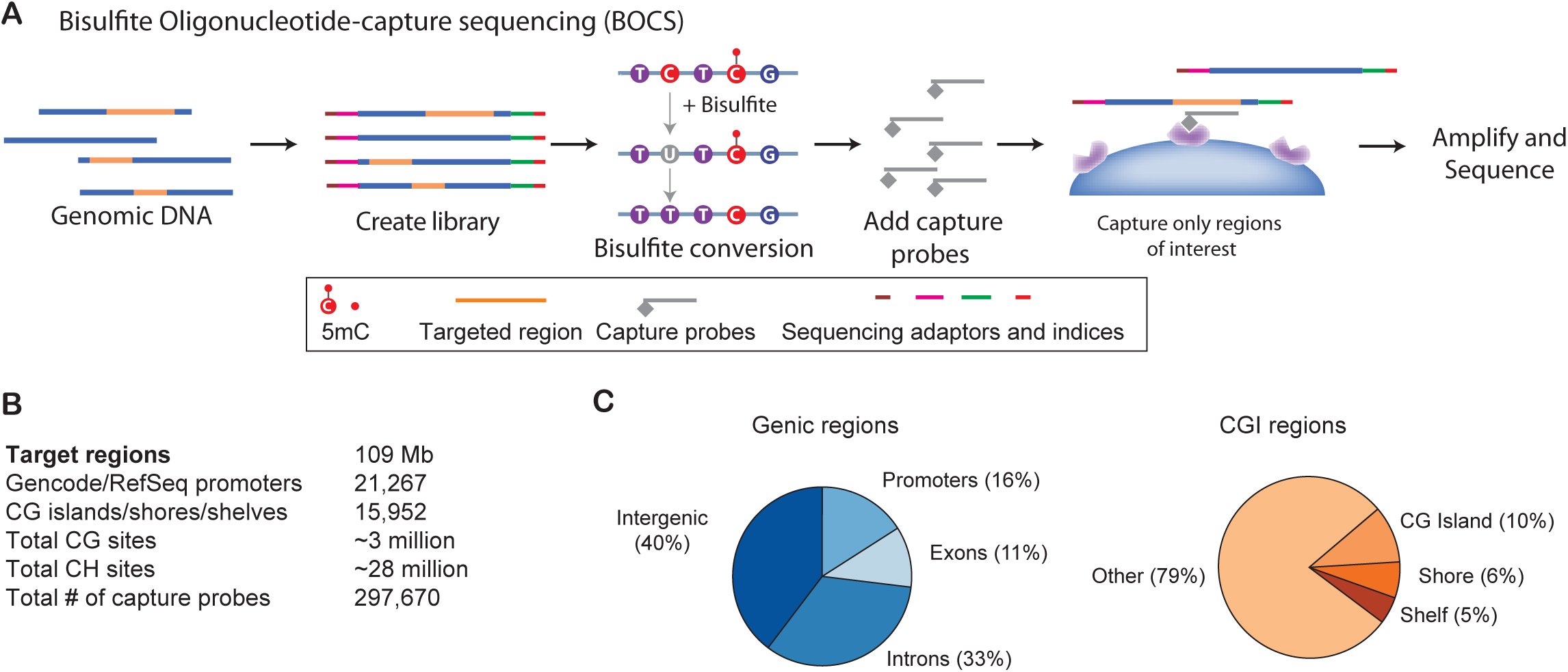
**Bisulfite Oligonucleotide Capture Sequencing (BOCS) method.** A) The BOCS methodology utilizes genomic DNA which is made into a sequencing library followed by bisulfite conversion. After conversion the target genomic regions of interest are captured with probes against methylated and unmethylated versions of the targeted genomic regions. The target regions are then captured and amplified prior to sequencing. B) Targeted regions of the mouse genome include 109Mb containing most of the annotated promoters and CpG islands. In total nearly 3 million CG and 28 million CH sites are in the targeted regions. C) Distributions of the targeted regions according to genic and CpG island context.

**Supplemental Figure 3:**
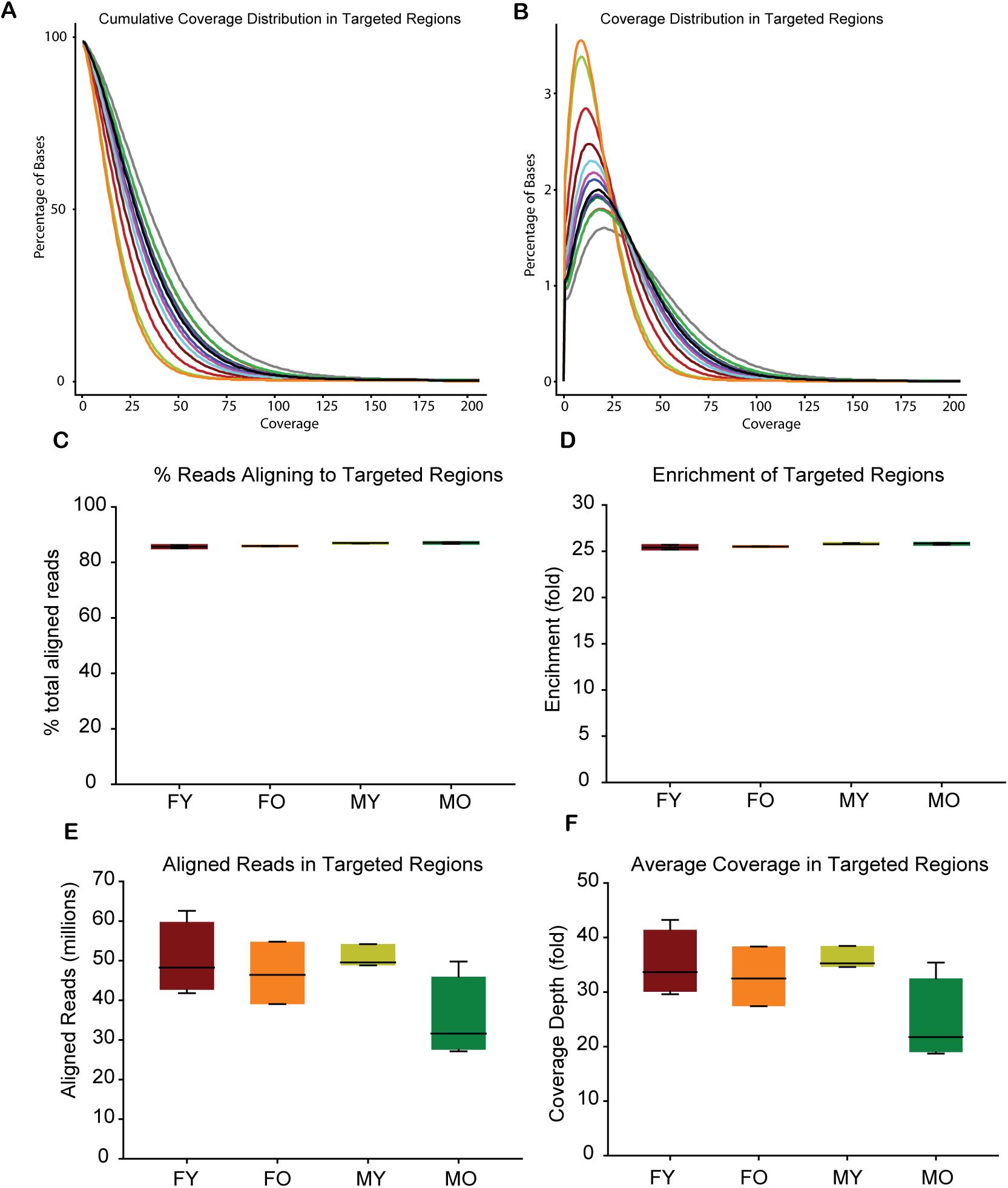
**Bisulfite oligonucleotide capture sequencing metrics.** After aligning reads to the full mm10 genome, for the portion of the genome targeted, a cumulative coverage distribution (A) and coverage distribution (B) for the base-level coverage depth in the targeted regions was generated. Each line represents an individual sample. C) A high percentage of aligned reads corresponded to the targeted regions demonstrating specificity of oligonucleotide capture. D) Average fold enrichment, coverage in targeted regions as opposed to the rest of the genome was greater than 25 fold in all groups. Total number of aligned reads (E) and average fold base coverage (F) for the targeted regions are also presented.

**Supplemental Figure 4:**
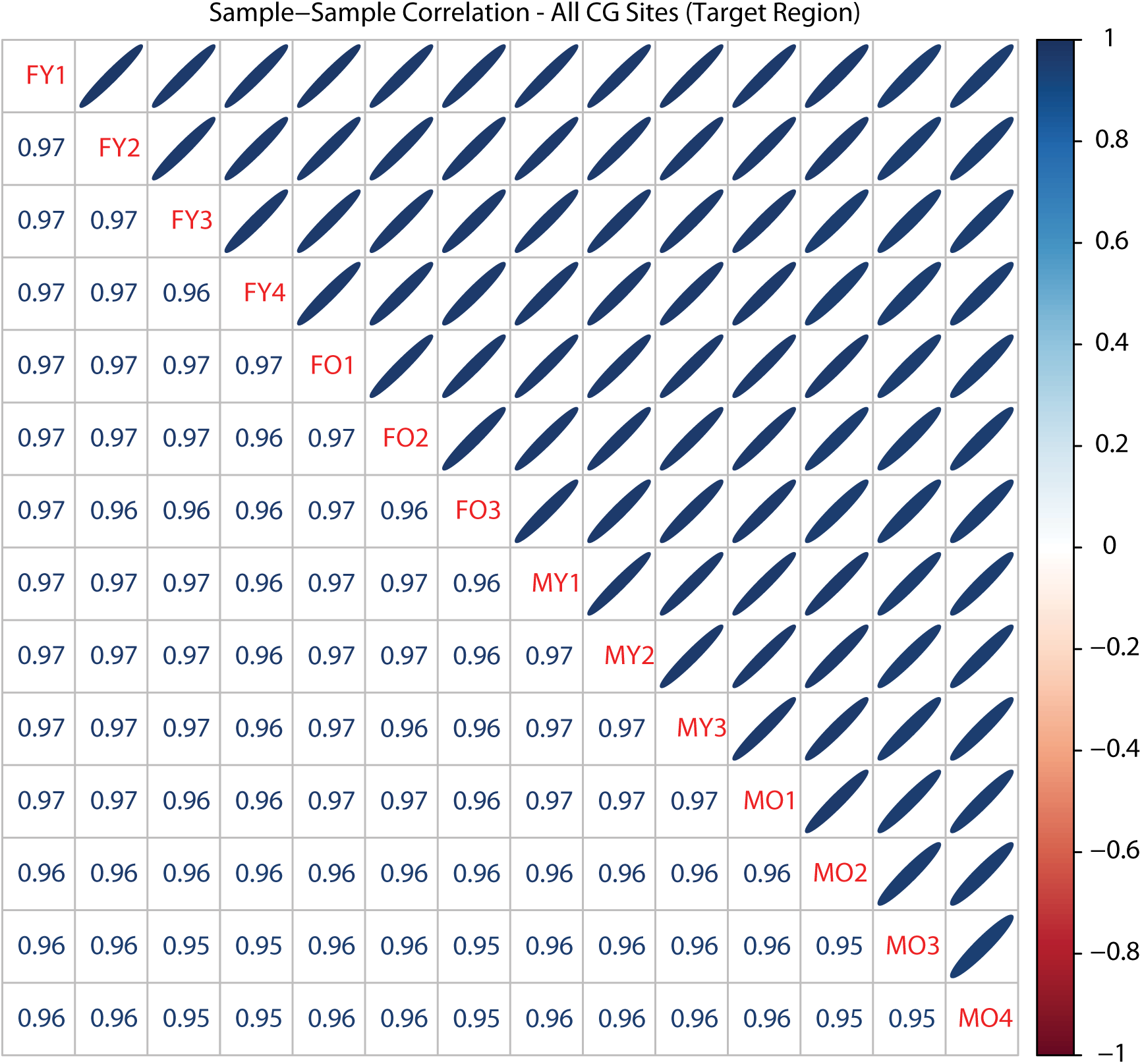
**Sample-sample correlations.** All pair-wise sample-sample correlations are presented for the full data set. All CG sites meeting coverage requirements were used in the correlation. All samples showeda correlation of 0.95 or higher demonstrating a high degree of technical reproducibility.

**Supplemental Figure 5:**
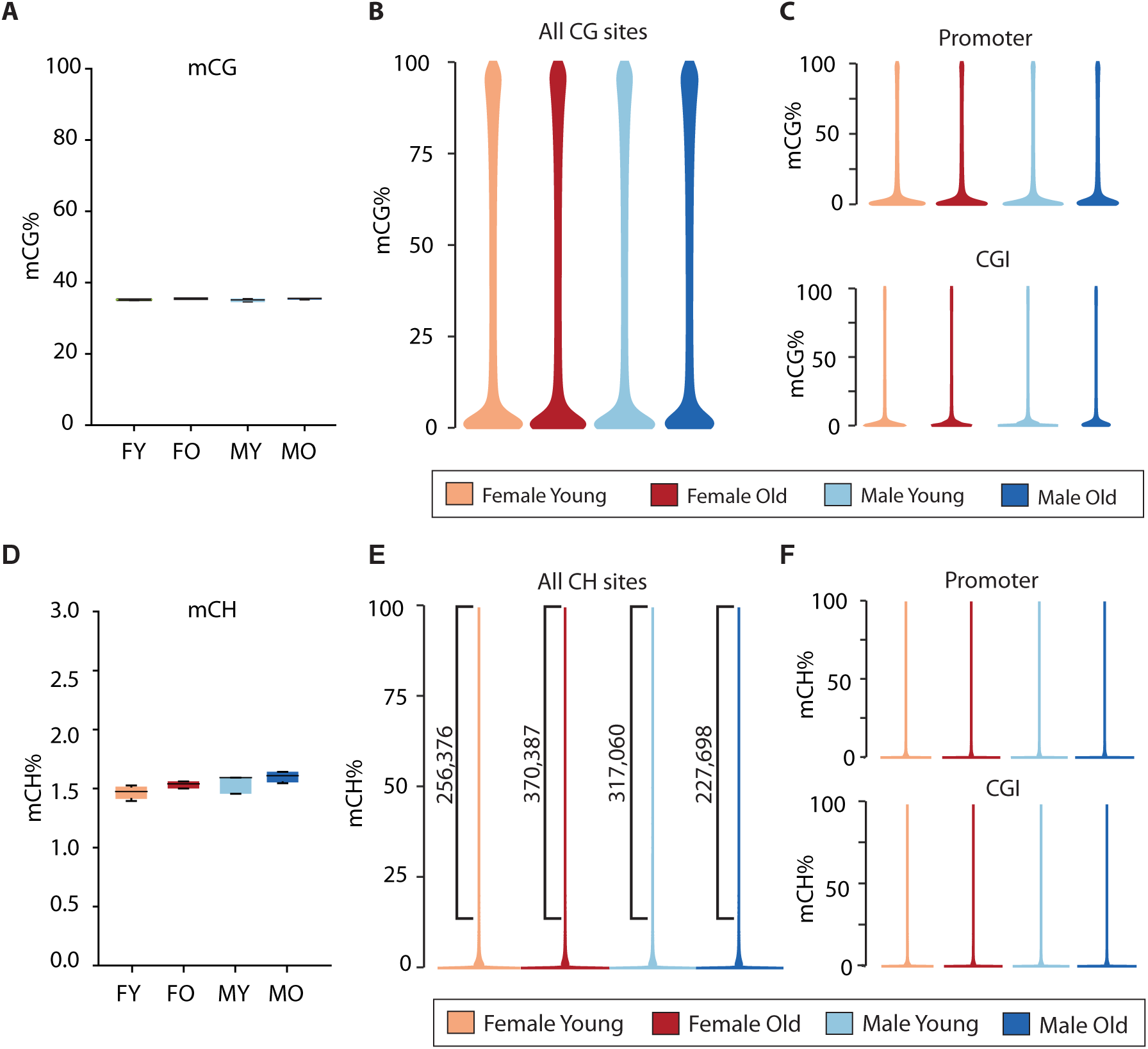
**Global mean methylation with age in female and male hippocampus.** A) Average CG methylation in young female (FY, 35.4%), old female (FO, 35.6%), young male (MY, 35.3%), and old male (MO, 35.3%) of all single site methylation calls (mean methylation at CG sites within targeted regions) demonstrated no differences with age or sex (Two-Way ANOVA with Age and Sex as factors, Student-Newman-Keuls post-hoc (SNK), Age p=0.216, Sex p=0.323, Sex and Age p=0.730, n=3-4/group, N=14). B) Density distribution of all CG sites genome-wide, in promoters C), and in CG Islands. D) CH (H is either A, C, or T) methylation levels in FY, FO, MY, and MO were equivalent across all groups and demonstrated no differences by age or sex. E) Density distribution of CH methylation genome-wide. Brackets represent the number of CH sites from the portion of distribution in each group (FY – 256,376, FO – 370,387, MY 317,060, MO – 227,698) that have CH methylation ≥ 10%. F) CH site density distributions across young and old males and females in promoter (top) and CG Island (bottom) regions.

File S1.xlsx, Age-related methylation differences at CG sites and CH sites

File S2.xlsx Autosomal life-long sex differences in CG and CH sites.

## Supplemental Methods

### Animals

Estrous cycle staging was performed by daily vaginal lavage to control for cycling differences. Lavages were conducted as described previously (Mangold *et al.* 2017) and using well established methods (McLean *et al.* 2012) to prevent any unnecessary stress on the animals. Briefly, sterile filtered water was expelled and aspirated approximately 4-5 times into the vaginal canal until enough cells were obtained for cytological analysis. Water from the vaginal wash was then placed onto a glass slide, allowed to dry, then stained using 0.1% crystal violet. The estrous cycle consists of 3 major phases: proestrus (high estrogen), estrus (low estrogen), and diestrus (low estrogen). Proestrus is defined by having a predominance of round, nucleated epithelial cells, estrus by cornified squamous epithelial cells, and diestrus by leukocytes with few epithelial cells present (McLean *et al.* 2012).

### Bisulfite oligonucleotide-capture sequencing (BOCS)

Genomic DNA (gDNA) was isolated from flash hippocampal frozen tissue using silica spin-columns (Zymo Duet) as described previously (Masser *et al.* 2016; Mangold *et al.* 2017). gDNA was quantified by fluorescent assay (PicoGreen, Invitrogen). 3 µg of gDNA for each sample was brought up to 50 µl volume with 1X TE and sheared by sonication (Covaris S2) to an average basepair size of 160 using the following settings; intensity of 5, duty cycle of 10%, 200 cycles per burst, 6 cycles of 60 seconds, at 4 °C. The size of sheared products was confirmed by capillary electrophoresis (DNA 1000, Agilent). gDNA libraries (SureSelect XT Methyl-Seq) were then constructed (Additional file 1: Figure S1A) following the manufacturer’s instructions (Agilent). gDNA fragments were end-repaired in 1X XT2 End-Repair Master Mix, and incubated for 30 minutes at 20 °C. End-repaired gDNA fragments were cleaned by magnetic SPRI-bead method (AMPure XP, Beckman-Coulter) and confirmed to be between 140 and 180 basepairs by capillary electrophoresis (DNA 1000). Fragments were then 3’ adenylated in 1X XT2 dA-Tailing Master Mix and incubated for 30 minutes at 37 °C. Methylated adapters were ligated on the gDNA fragments in XT2 Ligation Master Mix, incubated for 15 minutes at 20 °C, cleaned, and confirmed to be between 170 and 230 basepairs by capillary electrophoresis (DNA 1000). gDNA libraries were hybridized to the SureSelect Mouse Methyl-Seq Capture Library (Hing *et al.* 2015) (Agilent) for 24 hours at 65 °C. Capture baits were targeted for CGI units [CG Island ± 4kb (shores and shelves)], Gencode promoters, RefSeq promoters, and regulatory features including DNase I hypersensitivity sites and Ensembl regulatory features (Additional file 1: Figure S2B). Capture Library-hybridized DNA libraries were isolated and eluted using streptavidin beads. Captured DNA libraries were bisulfite converted using the EZ DNA Methylation-Gold method (Zymo Research), PCR amplified for 8 cycles, and bead purified. Libraries were indexed by PCR for 6 cycles, purified, and confirmed to be a library size of approximately 220 bp (High Sensitivity DNA Chip, Agilent). One old female and one young male library failed QC and were not included in the analysis. Libraries were quantified using standard curve qPCR (KAPA Biosystems), diluted to 4nM and pooled at equal volumes. Pooled libraries (12pM) were sequenced using the Illumina HiSeq 2500 and 2×150 Paired-End Rapid Run. All raw fastq files are publicly available from the NCBI Sequence Read Archive with the accession number PRJNA290881.

### Bioinformatics

Prior to alignment paired-end reads were adapter trimmed and filtered in CLC Genomics Workbench 8.5.1., while end-trimming removed 3 bp and 4 bp from the 5’ and 3’ end of each paired-end read, respectively. Only reads with a Q-score >30 were used for mapping, reads which did not meet criteria after trimming were discarded. Alignment of trimmed bisulfite converted sequences was carried out using Bismark Bisulfite Mapper v0.14.4 (Krueger & Andrews 2011) against the mouse reference genome (GRCm38/mm10). Our data analysis pipeline included samtools (Li *et al.* 2009), bedtools (Quinlan & Hall 2010), and R scripts while analysis of methylation was primarily done in R v3.3.0 using the package MethylKit (Akalin *et al.* 2012).

Total mean methylation levels for CGs and CHs across the whole genome by counts were calculated averaging all per site methylation calls (CGs and CHs separately) which were determined by dividing the methylated counts over total read counts for each site. The mean methylation fraction was then multiplied by 100 (mean methylation call = (methylated counts /total read counts) × 100). Only cytosines in the BOCS targeted regions, covered by at least 10 reads, and that were present in all samples within a comparison were included for statistical analysis of age changes and sex differences, resulting in 908,118 CGs and 13,423,219 CHs analyzed for differential methylation.

Differentially methylated cytosines (DMCG and DMCH) were determined from those sites passing coverage criteria. Differentially methylated sites between groups were determined using logistic regression (Akalin *et al.* 2012). P-values were adjusted using a false-discovery rate, or FDR (Benjamini & Hochberg 1995). Only sites with a q-value < 0.05 and an absolute methylation difference of ≥ 5% were considered significant. Sex differences were limited to those sites with coordinate differential methylation at both young and old ages.

Differentially methylated sites were examined for enrichment in specific genomic elements and annotated using the R package GenomicFeatures. Refseq genes and CGI unit’s coordinates were downloaded from UCSC genome browser (GRCm38/mm10) (https://genome.ucsc.edu/). Sites were examined for overlap with genic regions: introns, exons and promoters (defined as 3kb upstream and 300bp downstream of the transcription start site) or CGI units [CG Island including shores (± 2kb from islands), and shelves (± 2kb from shores), or CG Island ± 4kb]. Regions to which no overlap with any Refseq annotated genes was found were determined to be present in intergenic regions. Regions with no overlap with CGI units were annotated as ‘other’. Two-tailed χ^2^ test (alpha = 0.05) was used to determine the statistically significant differences in observed versus expected genomic annotation frequencies of differential methylated sites. Data is presented as fractions related to the total distribution of the sites analyzed.

Epigenomic enrichment analysis was performed using GenomeRunner as described previously (Dozmorov *et al.* 2016). Briefly, the enrichment analysis evaluates whether sites co-localizes with genome annotation datasets in a statistically significant manner. As the NCBI37/mm9 mouse genome assembly remains the best source of genome annotation datasets, the enrichment analysis was performed using mm9 genomic coordinates of ENCODE data from the UCSC genome browser database (Karolchik *et al.* 2014), accessed 07-22-2015 and a liftover from mm10 to mm9 for the DMCGs and DMCHs. The two-tailed χ^2^ test was used to calculate enrichment/depletion p-values, corrected for multiple testing using False Discovery Rate (FDR) approach, and odds ratios (limiting to only odds ratios <0.9 and >1.1).

Human brain methylation data were obtained from NCBI Gene Expression Omnibus (GEO), specifically control (not known to be diseased) samples from series GSE89703 (Viana *et al.* 2017) and GSE63347 (Horvath *et al.* 2015) for hippocampal data (22 samples, ages 25-95 years) and GSE41826 (Guintivano *et al.* 2013) for frontal cortex data (145 samples, ages 13-79). All data were generated from Illumina 450k Methylation Arrays and deposited onto GEO as beta values. Beta values and sample annotations were collected from GEO and no normalization or corrections were applied; rows with any missing values (NAs) and all sex chromosome sites were removed from analyses leaving 413085 probes for analysis in hippocampal samples and 469680 probes for analysis in frontal cortex. A set of general linear models (Ritchie *et al.* 2015) were used to observe the effects of age on methylation in hippocampus and frontal cortex (Figure 5 Panel A) and to test the main effects of age on methylation and sex as well as a model of the sex by age interaction for the frontal cortex (limiting to 69 male and 70) females samples ranging from 13-60 years of age, Figure 5 Panel B-E). Multiple testing correction was done for false discovery rate and a corrected p value of less than 0.05 was considered significant. All statistics were done using R statistics platform (Team 2015) and plotted using ggplot2 (Wickham 2009) and associated themes (Arnold 2017).

